# A complex three-dimensional microfluidic model that mimics the early stage events in the human atherosclerotic artery

**DOI:** 10.1101/2023.02.02.526873

**Authors:** Ranganath Maringanti, Christian G.M. van Dijk, Elana M. Meijer, Maarten M. Brandt, Merle M. Krebber, Ihsan Chrifi, Dirk J. Duncker, Marianne C. Verhaar, Caroline Cheng

**Author notes:** **Corresponding author:** Caroline Cheng, PhD, University Medical Center Utrecht, PO Box 85500, 3508 GA Utrecht, The Netherlands. These authors contributed equally to the manuscript (Christian G.M. van Dijk, Elana M. Meijer).

## Abstract

**Background:** Atherosclerosis is a complex inflammatory vascular disease characterized by lipid and immune cells accumulation in the vessel wall, leading to lumen narrowing. Although several 3D *in vitro* microfluidic systems were previously described, a realistic reconstruction of the *in vivo* human atherosclerotic environment requires co-culture of different cell types arranged in atherosclerotic vessel-like structures with exposure to flow and circulating cells, creating challenges for disease modelling.

In this study we developed a 3D tubular microfluidic model with quadruple coculture of human aortic smooth muscle cells (hAoSMCs), human umbilical cord vein endothelial cells (HUVECs) and foam cells to re-create a complex human atherosclerotic vessel *in vitro* to study the effect of flow and circulating immune cells.

**Methods & Results:** Our new co-culture protocol with BFP-labelled hAoSMCs, GFP-labelled HUVECs and THP-1 macrophages-derived, Dil-labelled Oxidized Low-Density Lipoprotein (Dil-Ox-LDL) foam cells in a fibrinogen-collagen-I based 3D extracellular matrix (ECM) resulted in vessels with an early lesion morphology, showing a layered vessel-like composition with an endothelium and media, with foam cells accumulating in the sub-endothelial space. Perfusion for 24 hours of atherosclerotic and “healthy” vessels (BFP hAoSMCs and GFP HUVECs without foam cells) showed that the layered wall composition remained stable. Perfusion with circulating THP-1 monocytes demonstrated cell extravasation into the atherosclerotic vessel wall and recruitment of THP-1 cells to the foam cell core. QPCR analysis revealed increased expression of atherosclerosis markers in the atherosclerotic vessels and adaptation in VSMCs migration to flow and the plaque microenvironment, compared to control vessels.

**Conclusion:** We present a 3D tubular microfluidic model of a complex early atherosclerotic human vessel that can be exposed to flow and circulating THP-1 monocytes to study hemodynamic changes and immune cell recruitment under live confocal imaging. This novel atherosclerosis-on-a-chip model offers a humanized platform for in-depth mechanistic *in vitro* studies and drug testing.

## Introduction

Atherosclerosis is a chronic inflammatory disease of the arteries, marked by the accumulation of cholesterol-containing low-density lipoproteins (LDL) leading to plaque formation and lumen narrowing^1^. As a leading cause of disease related death worldwide^2^, the aetiology of atherosclerosis is widely studied and is hallmarked by several causal factors including dyslipidaemia with high blood LDL levels, resulting in chronic inflammation and foam cell accumulation in the arterial wall. On a cellular level, it involves the activation of disease pathways in several vascular and immune cell types that interact with each other to amplify inflammation while adapting to (changes in) local hemodynamical factors. Traditionally, rodent disease models are used to provide valuable insights into the disease mechanism, but these are also known to diverge on multiple immune and mechanical aspects from the human condition, imposing limitations to their translational value^3–5^. Two-dimensional (2D) and three-dimensional (3D) human cell culture systems are also often used, for example to study vascular and immune cell responses to LDL and flow dynamics (reviewed in^6^). The most advanced of these 3D tools are based on tissue engineered (TE) vascular constructs, which can show a higher degree of complexity to mimic certain aspects of the atherosclerotic vessel, such as co-culture of multiple (vascular) cell types and organization of a layered wall structure^7–9^. Suitable for mechanistic studies and drug and toxicity testing in a controlled environment, they are cultured in static conditions^7, 8^, or exposed to lumen flow by perfusion in macroscale bioreactors when combined with mechanical support provided by (synthetic) scaffolds^9^ or hydrogels^10^. In recent years, microfluidic technology has allowed the introduction of luminal perfusion in complex microtissues. This adds critical flow dynamics (i.e. shear stress) to these biological models and bridges the gap between perfused culture with large (centimetre scale) bioreactor-based platforms to smaller, (micrometre scale) chip-format assays that can be more cost-effective as a result of their suitability for upscaling^11, 12^. In vascular research, multiple static models and perfused microfluidic devices have been developed to study endothelial function, angiogenesis, vasculogenesis, and inflammation^13–15^, but so far, only a handful of studies have presented an *in vitro* human cell microfluidic system for atherosclerosis^6^. In general, these studies demonstrated assays with co-cultures of vascular cells with immune cells that are organized in vascular-like structures, yet still lacked critical factors of atherogenesis, including physiologically relevant vessel anatomy and flow dynamics and vessel wall interaction with circulating immune cells^16–19^.

In this study, we created a complex, perfused human atherosclerosis on a chip (OAC) model that mirrors the atherosclerotic wall within a tubular arterial geometry in a full 3D extra cellular matrix (ECM) environment that allows physiological laminar flow patterns and natural interaction with circulating immune cells. This atherosclerosis model is based on a chip design that was recently published by our group^20^ and involves a 5 micro-channel system which can be further scaled up for the creation of multiple atherosclerotic vessels per chip for high throughput screening. Co-culture of endothelial cells, vascular smooth muscle cells (VSMCs), and (oxLDL loaded) foam cells in this full 3D ECM environment recreates a layered lesion structure that mimics the human atherosclerotic lesion in the initial phase of the disease. Fluorescent labelling of each individual cell type allows spatiotemporal visualization of cell behaviour during the different stages of immune cell-lesion interaction under live confocal imaging under luminal flow.

This new complex human atherosclerosis-OAC model is suitable for mechanistic studies of atherogenesis and drug testing that focusses on the contribution of circulating factors and immune cells on plaque progression. The system may also be further adapted to incorporate patient-derived cells (of induced pluripotent stem cell (iPSC) and/or PBMC origin), to create dedicated platforms for drug screening for patient populations with a genetic predisposition to coronary artery disease (CAD).

## Materials and Methods

Detailed descriptions can be found in the supplemented data.

### Cell culture (2D)

#### Primary cell culture

Human aortic smooth muscle cells (hAoSMC, Lonza) with a blue fluorescent protein (BFP)-tag were cultured in smooth muscle cell growth medium-2 (SMGM2) (Lonza). Human umbilical cord vein endothelial cells (HUVEC; Lonza) with a green fluorescent protein (GFP)-tag were cultured in endothelial growth medium-2 (EGM2).

#### Cell line /Suspension culture

THP-1 monocytes (ATCC TIB-202^TM^) were cultured in RPMI 1640 medium + L Glutamine (Life Technologies, Grand Island, NY) supplemented with 10% FBS (Gibco).

#### Differentiation of THP-1 monocytes into macrophages and foam cells

THP-1 cell differentiation into macrophages was induced by phorbol myristate acetate (PMA) in 10% FCS/RPMI medium. Macrophages were differentiated into foam cells by treating with Dil-labelled Ox-LDL (Invitrogen) with a concentration of 20 µg/mL for 24 h in RPMI bare medium.

### Cell culture (3D)

#### A) Co-culturing of vascular cells in a 3D microfluidic system

BFP-labelled hAoSMCs at a 10×10^6^ cells/mL concentration were seeded into the channels on day 1. The bioreactor was placed in 30 mm dish and rotated for 1-2 h on a MACSmix^TM^ tube rotator (Miltenyl biotech) for uniform spreading and attachment of cells in the channel. After rotation, bioreactors were submerged in SMGM2 medium and incubated. 12×10^6^ cells/mL GFP-labelled HUVECs were seeded in the same channels on day 2 and 3 and submerged in EGM2 and SMGM2 medium (ratio 1:1).

#### B) Labelling of monocytes or foam cells

THP-1 monocytes or foam cell suspensions were incubated with 1 µM Cell-tracker deep red solution for 30-45 min.

#### C) Co-culturing of circulating immune cells in a 3D microfluidic system

Cell-tracker deep red labelled THP-1 monocytes were perfused through the EC-VSMC co-cultured channels for 24 h in a cocktail medium (SMGM2: EGM2: RPMI-1:1:1) via the Ibidi pump system (Ibidi, Germany), at 5×10^5^ cells/mL circulating medium.

#### D) Co-culturing of vascular cells and foam cells in a 3D microfluidic system

To mimic atherosclerosis in the 3D microfluidic system, BFP-hAoSMCs were co-cultured with foam cells (pre-differentiated in 2D) and seeded in the channels. Foam cells and BFP-hAoSMCs were seeded in the ratio of 1:400, with foam cells and BFPAoSMCs concentrated to 25000 and 10 million cells/mL respectively. To monitor if only macrophages have taken up Dil-ox-LDL in the experiments presented in figure 4 and 5, foam cells were also stained with cell-tracker deep red before seeding. After seeding into the channels, the bioreactors were rotated and incubated followed by seeding of GFP-HUVECs on day 2 and 3. Finally, the bioreactors were incubated in cocktail medium (SMGM2:EGM2: RPMI, ratio 1:1:1).

#### Perfusion experiments (3D)

After 4 days of static co-culture, the microfluidic devices were subjected to flow using the Ibidi pump system. A selection of channels was connected to the perfusion system via 26G needles (perfused group). The others were closed with PE-50 tubing (static group). To allow the cells to adjust to unidirectional flow, 20 µl/min flow rate was first maintained for one hour before it was increased to 40 µl/min.

#### Confocal Imaging

Imaging was performed by Leica SP8 confocal microscope using 10x magnification for both z-stack mode (12 μm step size) and tile scan mode (3×2). Image analysis was performed with Leica Application Suite X software, (version 3.7.1.21655) and ImageJ.

#### Gel excision from 3D microfluidic device and qPCR analysis

Following confocal readout, the microfluidic devices were snap frozen on dry ice and the gel was excised from the bioreactor and dissolved in RNA lysis buffer. Total RNA was isolated from the channels using RNA isolation kit (Bioline) for gene expression analysis.

#### Statistics

Statistical analysis was performed using GraphPad Prism 9.0. Data were represented as means ± SEM. Groups were compared using unpaired t-test or two-way or one-way ANOVA followed by Dunnett’s multiple comparison test or Tukey post hoc test, when appropriate. If other statistical methods were used, it was stated separately in the legends. Statistical significance was accepted when P ≤ 0.05.

## Results

### Medium optimization and creation of a human macrovessel mimic in a 3D microfluidic system for live confocal imaging

The human primary vascular cell types required for the creation of macrovessels were first assessed on cell survival and proliferation capacity in response to different growth medium conditions. hAoSMCs and HUVECs were cultured either in bare basal medium (SMBM2 bare, EBM2 bare or RPMI bare), their suitable full medium (SMGM2 or EGM2, for hAoSMCs and HUVECs respectively) and 2 cocktail media (SMGM2:EGM2 in 1:1 ratio or SMGM2:EGM2:RPMI in 1:1:1 ratio). The inclusion of RPMI in the second cocktail medium is for the final co-culture protocol with hAoSMCs, HUVECs and THP-1 cells. The cells were maintained for 7 days and cell viability was determined using PrestoBlue viability assay. Both hAoSMCs and HUVECs cultured in full growth medium SMGM2 or EGM2 (respectively), and cocktail medium SMGM2:EGM2:RPMI showed significant increase in cell viability compared to bare medium (SMBM2 bare or EBM2 bare, and RPMI bare). Cell culture of hAoSMCs or HUVECs in cocktail medium SMGM2:EGM2 or SMGM2:EGM2:RPMI showed comparable cell viability versus full growth mediums SMGM2 or EGM2 (Supplemental Fig. 1a & c). PicoGreen assay as a measurement for cell proliferation showed significant increase in DNA content in hAoSMCs and HUVECs cultured in their full growth medium SMGM2 and EGM2, and in the cocktail medium SMGM2:EGM2:RPMI versus their relevant bare medium conditions and RPMI bare.

hAoSMCs or HUVECs cultured in cocktail medium SMGM2:EGM2 or SMGM2:EGM2:RPMI showed comparable DNA levels versus their relevant full medium conditions (Supplemental Fig. 1b & d). These observations indicate that the media SMGM2:EGM2 and SMGM2:EGM2:RPMI are suitable for coculture of hAoSMCs and HUVECs in the 3D microfluidic system as cell viability and growth are not negatively affected. For the use of THP-1 cells as circulating monocytes and differentiated macrophages, we tested the viability of THP-1 cells in full medium (RPMI 10% FCS), bare medium (RPMI), and cocktail medium (SMGM2:EGM2:RPMI, ratio 1:1:1). Both cell viability and DNA content showed a significant higher cell viability for THP-1 cells cultured in full growth medium versus bare medium (Supplemental Fig.1e & f). THP-1 cell viability was significantly increased in cocktail (SMGM2:EGM2:RPMI) versus bare and full medium conditions (Supplemental Fig. 1e & f). These data demonstrate that SMGM2:EGM2:RPMI is a suitable growth medium for the co-culture of human HUVECs and VSMCs with THP-1 cells.

The human artery is composed of an inner endothelial monolayer surrounded by VSMCs in the media. To engineer a human macrovessel consisting of a confluent endothelium and sub-endothelial VSMCs layers that can be monitored live in the microfluidic device, 500 µm diameter channels were created by casting of a mix of fibrinogen (20 mg/mL in SMGM2) and collagen-I (2 mg/mL in SMGM2) ECM gel in the chamber, followed by seeding of vascular cells (as described in Supplemental Fig. 2a). After 4 days of static co-culture, confocal analysis showed a human macrovascular-like structure composed of a uniform endothelial monolayer formed by GFP-HUVECs (green) and subendothelial BFP-AoSMCs layers (blue) (Fig. 1a, shown as merged maximum projection images of the vessel wall). The 3D composite cross-sectional display (Fig. 1b), 3D reconstruction of half of the macrovessel (Fig. 1c), as well as the longitudinal (Fig. 1d) and orthogonal cross sections (Fig. 1e) revealed the layered vessel anatomy (GFP-HUVECs endothelium on top of BFP-AoSMC media) and showed the open lumen structure (∼500 µm in diameter) of these tissue-engineered human macrovessels.

**Figure 1.**
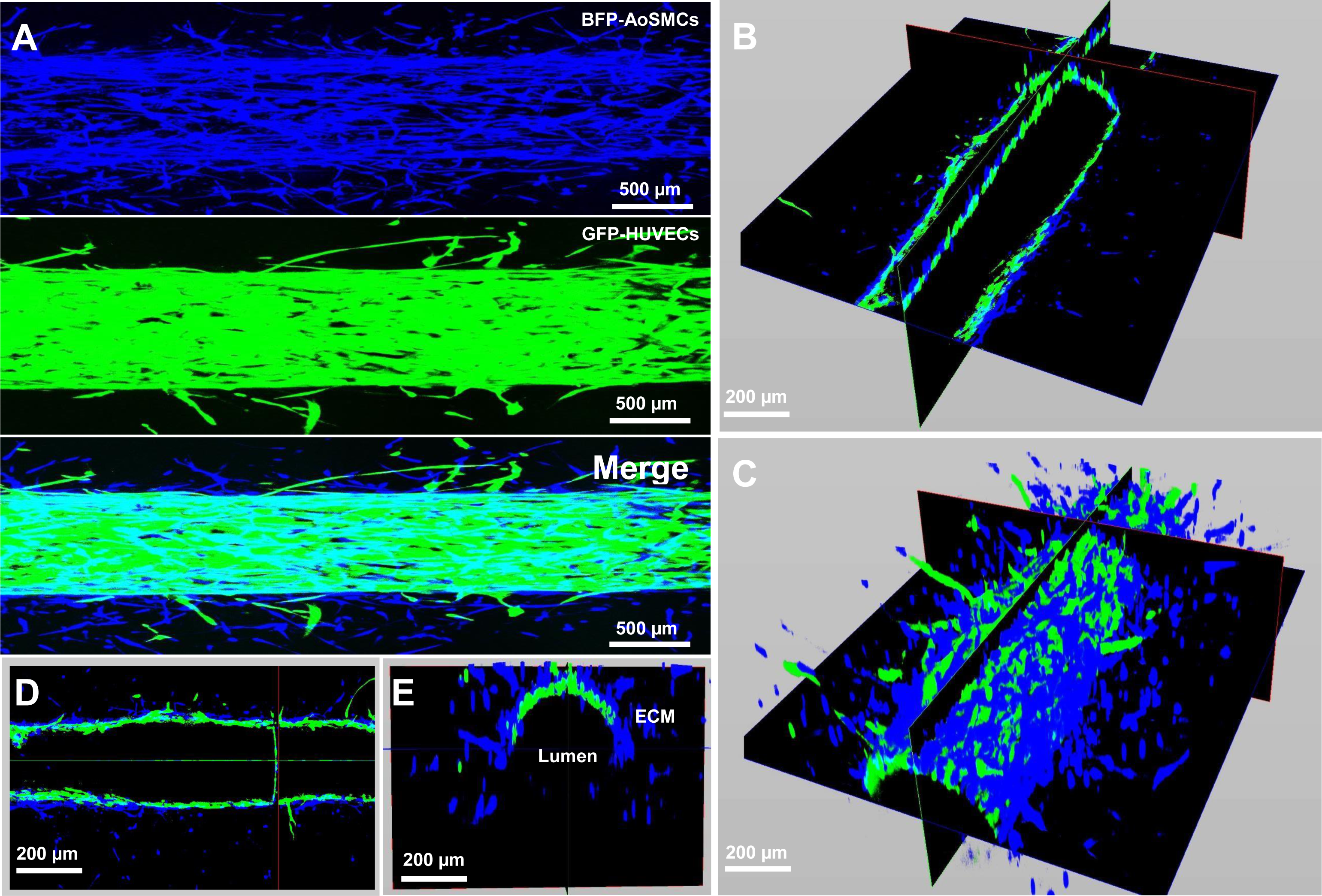
Fluorescent labeled human macrovessels mimicked in an in-house developed microfluidic device for live confocal imaging after 4 days of static coculture. (A) Confocal micrographs of vessel wall with an endothelial monolayer formed by GFP-labelled HUVECs (mid panel, in green) surrounded by BFP-labelled human AoSMCs (upper panel, in blue) and merged image (lower panel) of the created human macrovessel. (B) Composite display of the longitudinal and orthogonal cross section micrographs. (C) 3D reconstruction of half of the macrovessel wall. (D) Longitudinal cross section of the vessel. (E) Composite display of the orthogonal cross section micrographs showing the open lumen and layered structure of half of a macrovessel.

### Continuous vessel stability after introduction of flow in human macrovessels in the microfluidic system

Flow was introduced in the human macrovessels after 4 days of static co-culture by connecting the microfluidic device to a closed circulating Ibidi pump system. Cocktail medium (SMGM2:EGM2) was perfused unidirectionally through the macrovessel lumens at a flow rate of 20 µl/min for 1 h to allow the vessels to adapt from static condition to flow, before it was increased to 40 µl/min and maintained for 48 h (Supplemental Fig. 2b). This flow rate is within the range previously used in microfluidic platforms to study leukocyte–endothelium interaction under controlled flow^21–23^. Confocal imaging of the perfused channel after 48 h revealed preservation of the human macrovessel with an intact GFP-HUVECs (green) monolayer and subendothelial smooth muscle cell layers (BFP-hAoSMCs (blue)) surrounding the endothelium, composing the vessel wall (Fig. 2a). Evaluation by confocal 3D reconstruction (Fig. 2b and c) and analysis of the longitudinal (Fig. 2d) and orthogonal cross sections (Fig. 2e) further confirmed conservation of the endothelium and media and revealed full preservation of open lumen structures without noticeable changes in lumen diameter (Fig. 2 b-e). In line with these observations, quantification of the confocal images revealed that perfusion of the vessel for 48 h after 4 days of static coculture did not significantly affect the peri-luminal BFP-hAoSMCs+ and GFP-HUVECs+ areas when compared to 4 days static co-culture (Supplemental Fig. 4a and b).

**Figure 2.**
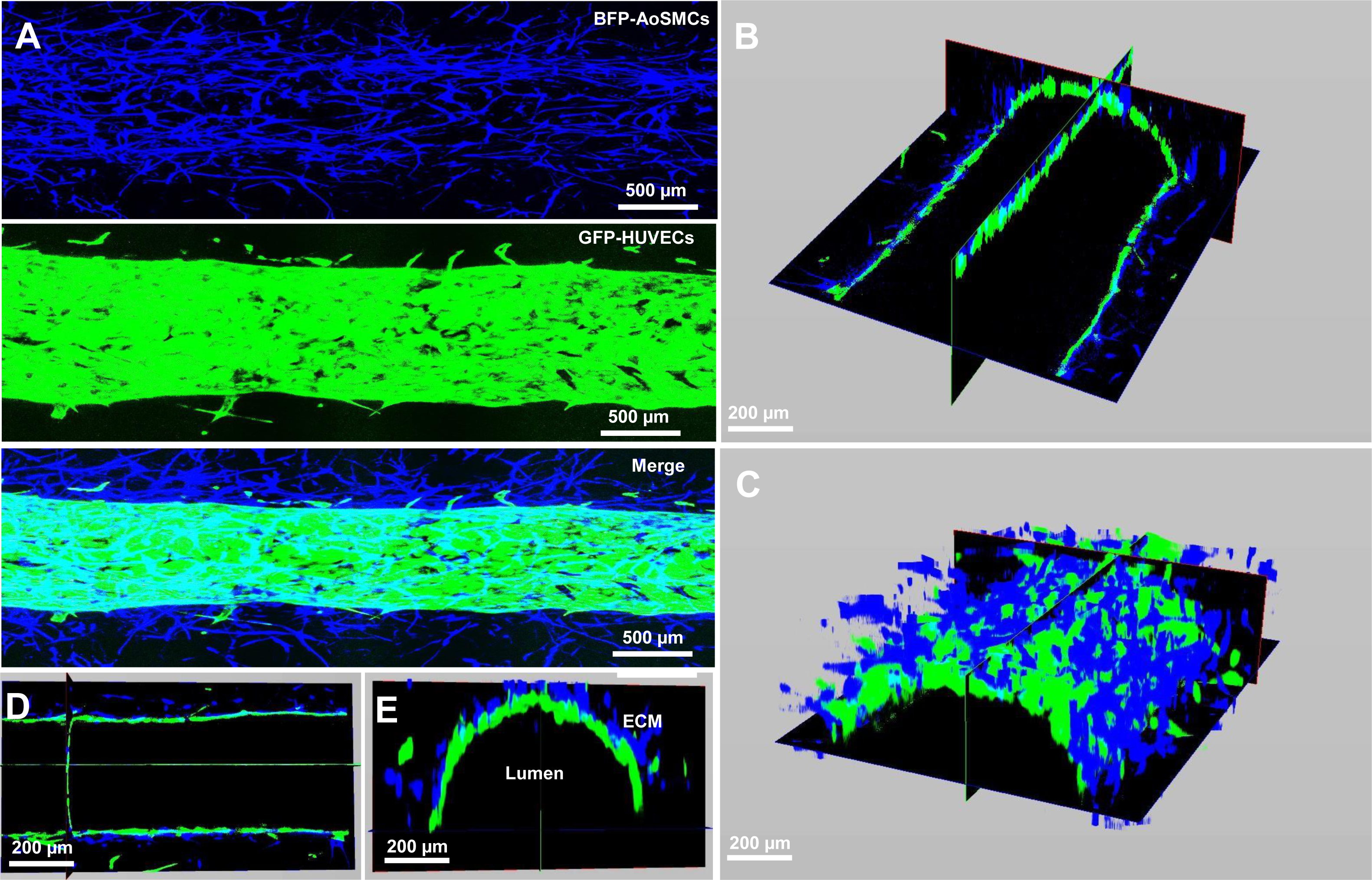
Human macrovessels after 48h of perfusion. (A) Confocal micrograph shows the preserved endothelial monolayer formed by GFP-HUVECs (mid panel, in green) surrounded by BFP-AoSMCs (upper panel, in blue), and merged image (lower panel) after 6 days of co-culture, from which 4 days static and 2 days perfused at a flow rate of 40µl/min. (B) Composite display of the longitudinal and orthogonal cross section micrographs. (C) 3D reconstruction of half of a wall of the perfused macrovessel. (D) Longitudinal cross section of the vessel. (E) Composite display of the orthogonal cross section micrographs showing half of a macrovessel, demonstrating preservation of the open lumen and layered structure.

### Preserved integrity after flowing with circulating monocytes in human macrovessels

In atherogenesis, monocyte recruitment and subsequent differentiation into macrophages and foam cells in the sub-endothelial space represent a hallmark process in disease onset and progression^24^. We introduced THP-1 monocytes that were labelled with cell tracker deep red in the cocktail medium and perfused these through the (healthy) control human macrovessels (Supplemental Fig. 2c) to assess THP-1 cell interaction with the vascular wall. Cell concentrations for flowing were optimized for THP-1 monocytes at 5×10^5^ cells/ml in SMGM2:EGM2:RPMI, at a flow rate of 40 µl/min. Confocal live imaging demonstrated limited THP-1 monocyte interaction with the control vessels (Fig. 3b-e). Confocal analysis after 48 h of continuous flow with THP-1 monocytes revealed preservation of the human macrovessels with intact lumen and endothelial and medial layers, similar to the perfused macrovessels without circulating THP-1 cells (Fig. 3a-e). Evaluation by confocal 3D reconstruction and analysis of the longitudinal and orthogonal cross sections further revealed that THP-1 monocytes (magenta) were mostly present in the lumen area (Fig. 3c-e, indicated by white arrows). A limited number of THP-1 localized at the adluminal surface of the GFP-HUVECs endothelial monolayer (Fig. 3b-e, indicated by yellow arrows), but none of these cells transmigrated deeper into the BFPhAoSMCs medial layer or further into the surrounding ECM. Similar to the flow condition without THP-1 cells, the BFP-hAoSMC^+^ and GFP-HUVEC^+^ areas were not affected by flowing with circulating THP-1 cells (Supplemental Fig. 4a and b).

**Figure 3.**
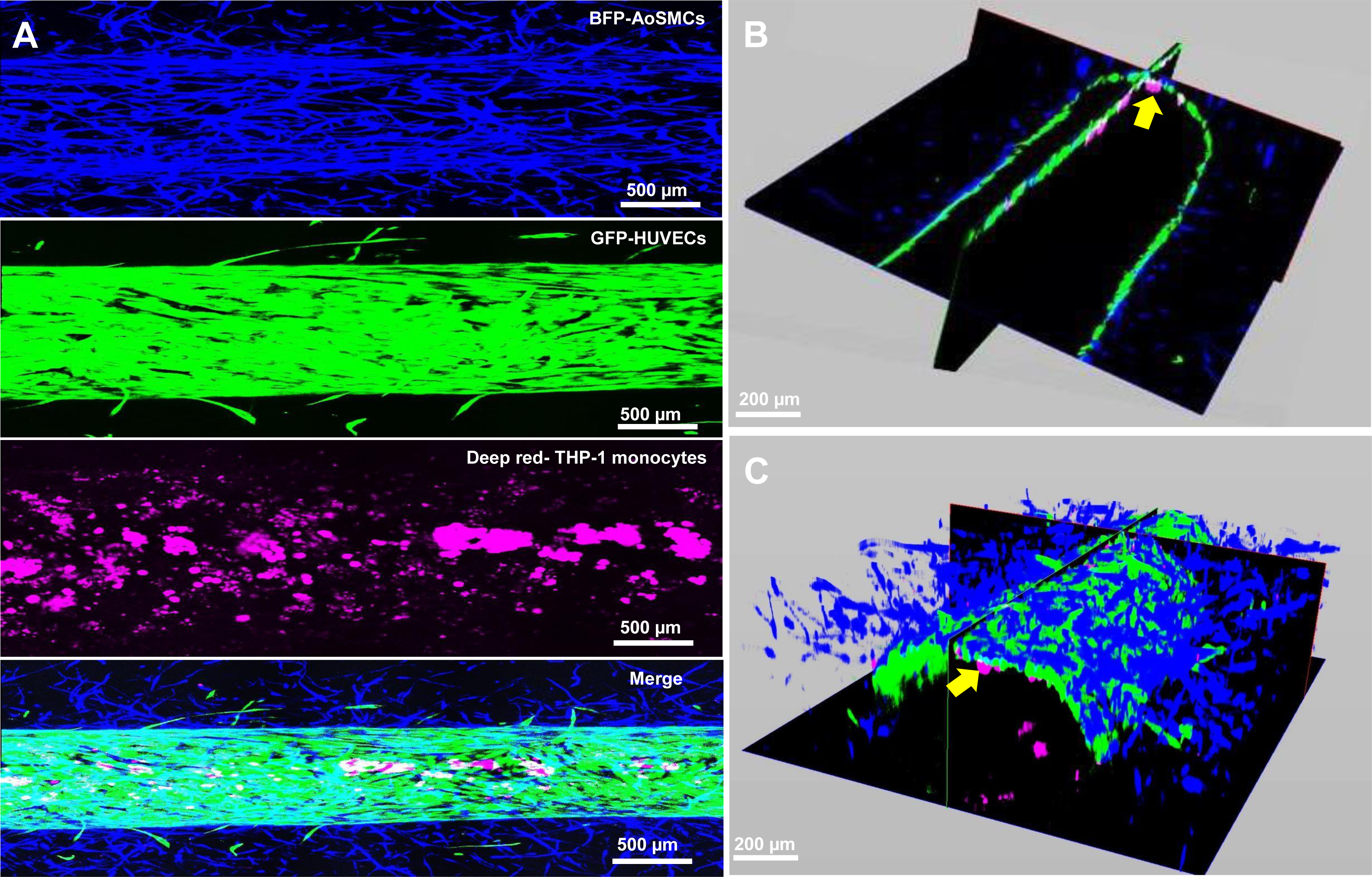
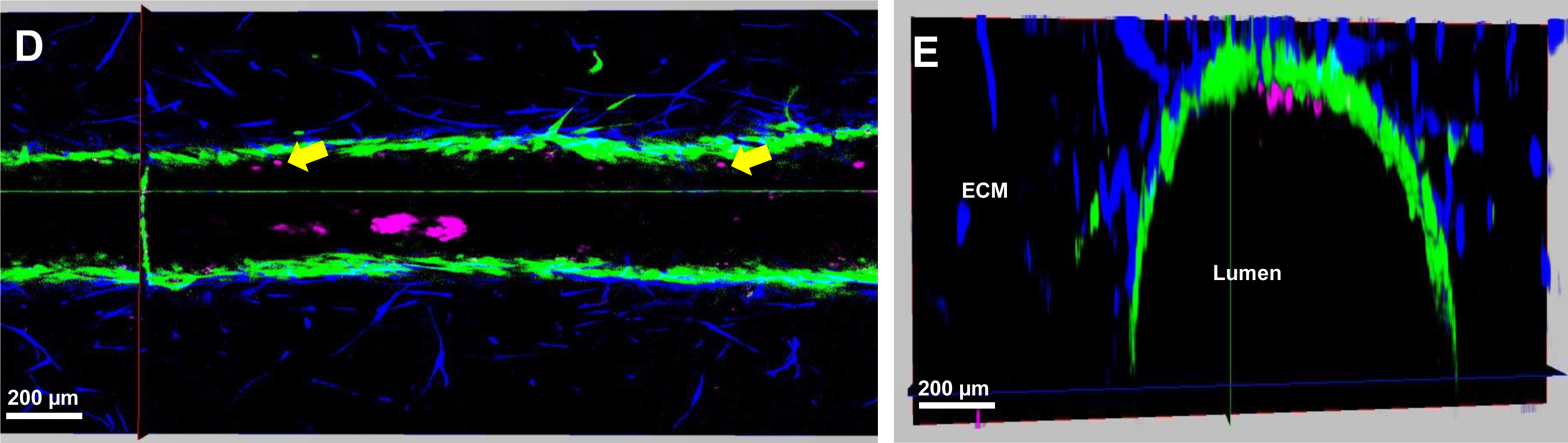
Human macrovessels after 48h of perfusion with circulating THP-1 monocytes. (A) Confocal micrograph shows the preserved endothelial monolayer formed by GFP-HUVECs (second panel, in green) surrounded by BFP-AoSMCs (first upper panel, in blue), as well as the deep red cell tracker stained THP-1 monocytes (third panel, in magenta) that were perfused in the human macrovessel after 6 days of co-culture, from which 4 days static and 2 days perfused with medium with circulating THP-1 cells at flow rate of 40µl/min. Lower panel shows merged image. (B) Composite display of the longitudinal and orthogonal cross section micrographs. C) 3D reconstruction of half of a wall of the perfused macrovessel. (D) Longitudinal cross section of the vessel. (E) Composite display of the orthogonal cross section micrographs showing half of a macrovessel, with preservation of the open lumen and layered structure. THP-1 cells remained mainly free-floating in flow medium and were localized in the lumen, although some of the THP-1 cells were interacting with the endothelium (yellow arrows).

### Optimization of Ox-LDL dose and introduction of foam cells in human macrovessels to create atherosclerotic macrovessels

Foam cell formation and accumulation in the sub-endothelial space of arteries is a hall mark of atherosclerotic lesions. We first determined the optimal oxidized-LDL (Ox-LDL) dose that is required for foam cells induction.

Under 2D culture conditions, THP1 monocytes were differentiated into macrophages using PMA and treated with Dil-labelled Ox-LDL for live fluorescent monitoring at a concentration of 20 µg/mL or 40 µg/mL for 24 h or 48 h. Ideal cell concentration for seeding was established at 5×10^4^ cells/mL of RPMI bare medium. Confocal imaging confirmed uptake of Ox-LDL (red) by macrophages at different doses and time points (Supplemental Fig. 3a). Quantification of the images showed a significant reduction in nuclei count at 40 µg/mL of Ox-LDL after 48 h of stimulation, whereas 20 µg/mL Ox-LDL did not impact cell numbers (Supplemental Fig. 3b). Similarly, quantification of the Dil-Ox-LDL signal showed that 20 µg/mL of Ox-LDL induced a higher number of Dil-Ox-LDL^+^ foam cells versus stimulation with 40 µg/mL (Supplemental Fig. 3c).

Cell viability and proliferation of THP-1 macrophages with and without exposure Dil-Ox-LDL was also evaluated by PrestoBlue and PicoGreen analysis. Similar to non-treated THP-1 macrophages, cell viability and proliferation declined over time (24 h versus 48 h) after 20 and 40 µg/mL Ox-LDL stimulation, although there was no impact of different Ox-LDL concentration on viability compared with the control macrophages (Supplemental Fig. 3d). The DNA content also declined over time for control macrophages, and macrophages stimulated with 20 and 40 µg/mL Ox-LDL. Ox-LDL stimulation (both 20 and 40 µg/mL) significantly enhanced the decline compared with control macrophages (Supplemental Fig. 3e). Based on these findings, we proceeded with 20 µg/mL 24 h Ox-LDL loading for foam cell induction.

Foam cells differentiated under 2D were harvested and stained with cell tracker deep red before seeding into the microfluidic system along with BFP-hAoSMCs on day 1 (at a concentration of 25000 foam cells to 10 million BFP-hAoSMCs -1:400) followed by GFP-HUVECs (Supplemental Fig. 2d). Confocal maximum projections at day 4 under static condition showed successful intra-wall foam cell accumulation (Fig. 4a, third panel in magenta) without negative impact on vessel integrity, shown by the general preservation of the endothelial monolayer (green) and VSMC layers (blue) (Fig. 4a, upper two panels). No significant differences were observed in the quantified BFPhAoSMCs^+^ and GFP-HUVECs^+^ areas between static co-culture with and without foam cell loading (Supplemental Fig. 4e and f). 3D reconstructions and cross sections demonstrate that an open lumen was also maintained (Fig. 4b-d). Higher magnification observation showed that most of the Magenta^+^ foam cells were located at the abluminal surface of the GFP-HUVECs endothelial monolayer with pockets of foam cells located deeper in the BFP-hAoSMCs medial layer further away from the endothelium (Fig. 4d and e, indicated by white arrows). Combined, this data indicates that this optimized protocol for foam cell introduction in “healthy” vessels, has successfully created a mimic of an early-stage atherosclerotic human vessel.

**Figure 4.**
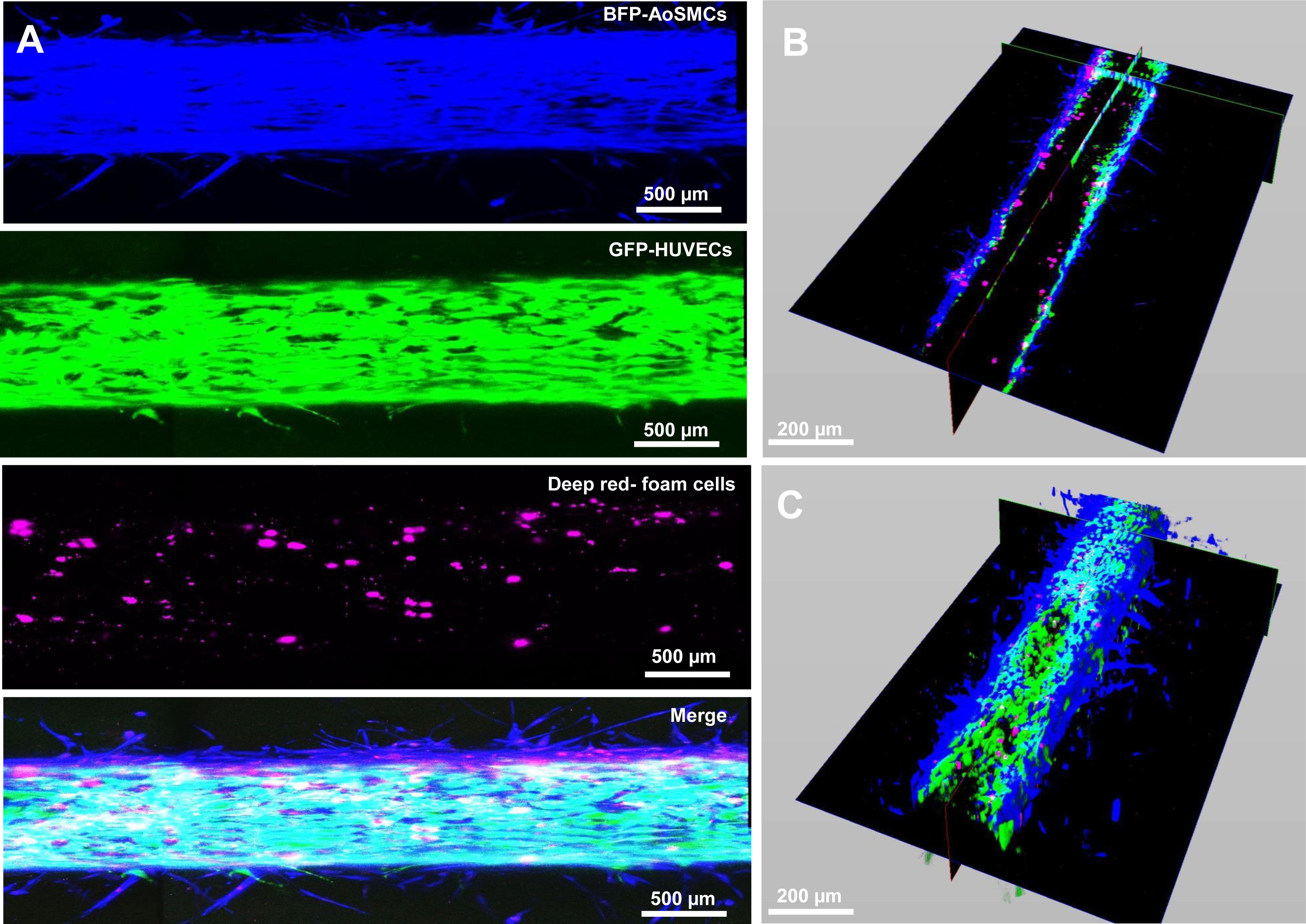
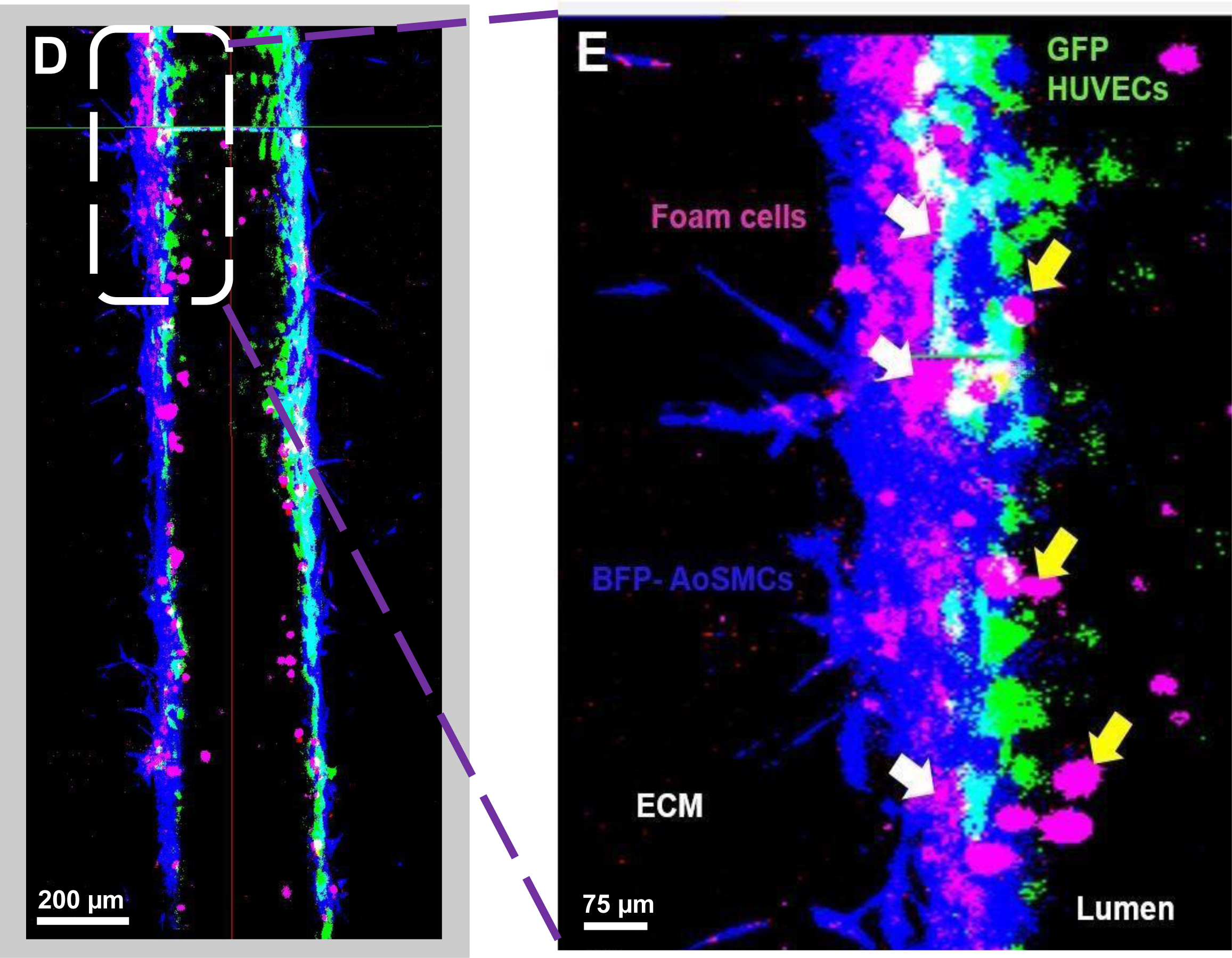
Introduction of foam cells in human macrovessels in the microfluidic system after 4 days of static co-culture. (A) Confocal micrographs of the preserved endothelial monolayer formed by GFP-labelled HUVECs (second panel, in green) surrounded by preserved BFP-labelled hAoSMCs (upper panel, in blue) co-cultured (static) with foam cells (third panel in magenta) and merged image (lower panel) of the created human atherosclerotic macrovessel. (B) Composite display of the longitudinal and orthogonal cross section micrographs. (C) 3D reconstruction of half of the macrovessel wall. (D) Longitudinal cross section of the vessel. (E) High magnification of a selected area in the longitudinal cross section showing most foam cells (magenta) accumulated in the smooth muscle cell layer (blue) just below the endothelium (green) (the sub-endothelial space (indicated by white arrows)) and some foam cells on the luminal surface (indicated by yellow arrows).

### Preserved vessel stability in perfused human atherosclerotic macrovessels in the microfluidic system

To allow studies of tissue adaptation to hemodynamic stimulation, the atherosclerotic vessels were subsequently tested with hemodynamic loading. Confocal live imaging following perfusion with the select cocktail medium for 24 h at a flow rate of 40 µl/min (Supplemental Fig. 2e) showed preservation of the vascular wall bilayer structure, and a conservation of the open lumen structure with the presence of foam cell accumulation in the sub-endothelial space, as shown by maximum projection images, 3D composite display, 3D reconstruction and longitudinal cross sections (Fig. 5 a-e). In line with these findings, quantification of the composite images showed no significant differences in BFP-hAoSMCs^+^ and GFP-HUVECs^+^ areas when compared to static atherosclerotic vessels (Supplemental Fig. 4c, d) or when compared to perfused control vessels (Supplemental Fig. 4g and h).

**Figure 5.**
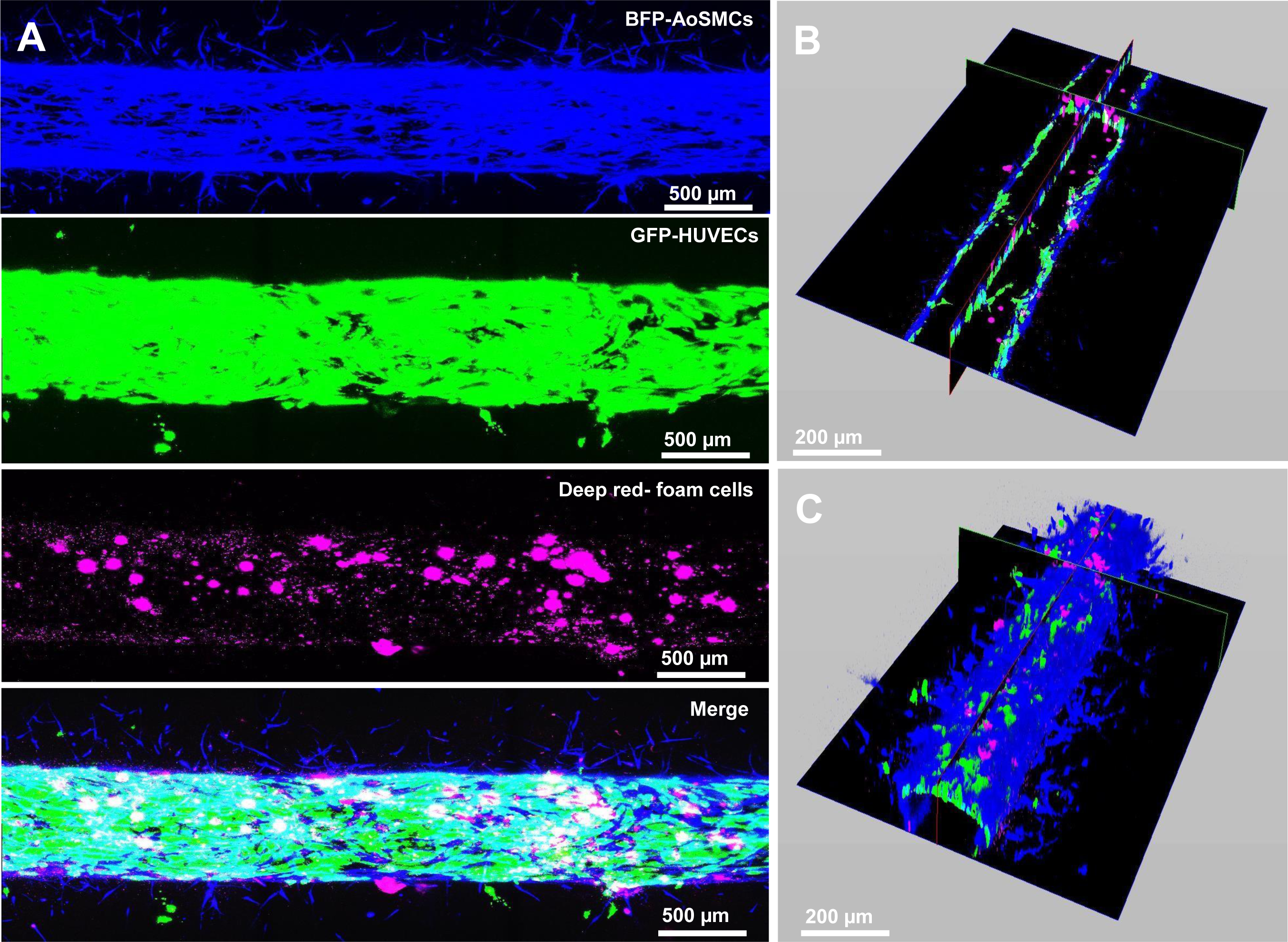
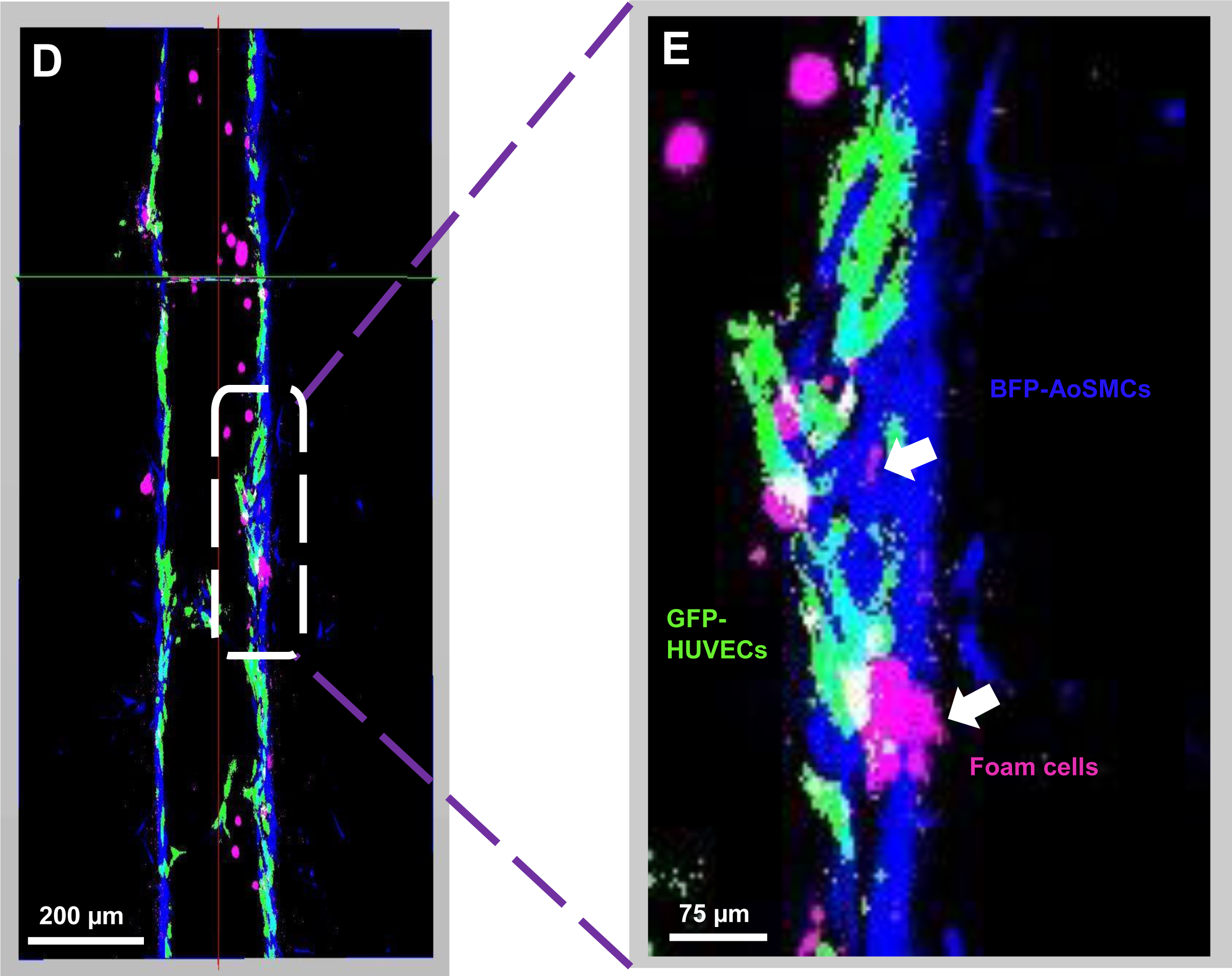
Flow perfusion for 24 hours in a human atherosclerotic vessel in the microfluidic system. (A) Confocal micrograph shows the preserved endothelial monolayer formed by GFP-HUVECs (second panel, in green) surrounded by preserved BFP-AoSMCs (upper panel, in blue) and accumulated foam cells (third panel in magenta), with the merged image (lower panel) after 5 days of co-culture, from which 4 days static and 1 day perfused at a flow rate of 40 µl/min. (B) Composite display of the longitudinal and orthogonal cross section micrographs. C) 3D reconstruction of half of a wall of the perfused macrovessel. (D) Longitudinal cross section of the vessel. (E) High magnification of a selected area in the longitudinal cross section showing foam cells (magenta) accumulation in the sub-endothelial space (indicated by white arrows) in the smooth muscle cell layer (blue) under the endothelium (green) as well as foam cells on the luminal side.

### Perfusion of human atherosclerotic vessels with circulating monocytes

Circulating monocytes extravasate into the sub-endothelium of the atherosclerotic plaque in response to local inflammatory signals, followed by macrophage differentiation and contribution to the foam cell core after Ox-LDL uptake during lesion progression. To develop the microfluidic human atherosclerotic chip model into a platform that can facilitate the study of this process, atherosclerotic vessels were perfused with cell tracker deep-red stained THP-1 monocytes, at a concentration of 5×10^5^ cells/ mL for 24 h at a flow rate of 40 µl/min (Supplemental Fig. 2f). Confocal live imaging demonstrated a high level of interaction between THP-1 cells and the vascular wall: Confocal maximum projections images showed distributions of GFP-HUVECs (green), BFP-AoSMCs (blue), Dil-oxLDL foam cells (red), and deep-red labelled circulating immune cells (magenta) in the vessel with general preservation of the atherosclerotic vascular wall anatomy (Fig. 6a). In the images of 3D composite display (Fig. 6b) and 3D reconstruction (Fig. 6c) infiltration of circulating monocytes (magenta) in the media was observed. High magnification display of a longitudinal cross-section with separated colour channels demonstrated significant THP-1 cells interaction with the endothelium, showing a high degree of THP-1 colocalization with HUVECs (Fig. 6d).

**Figure 6.**
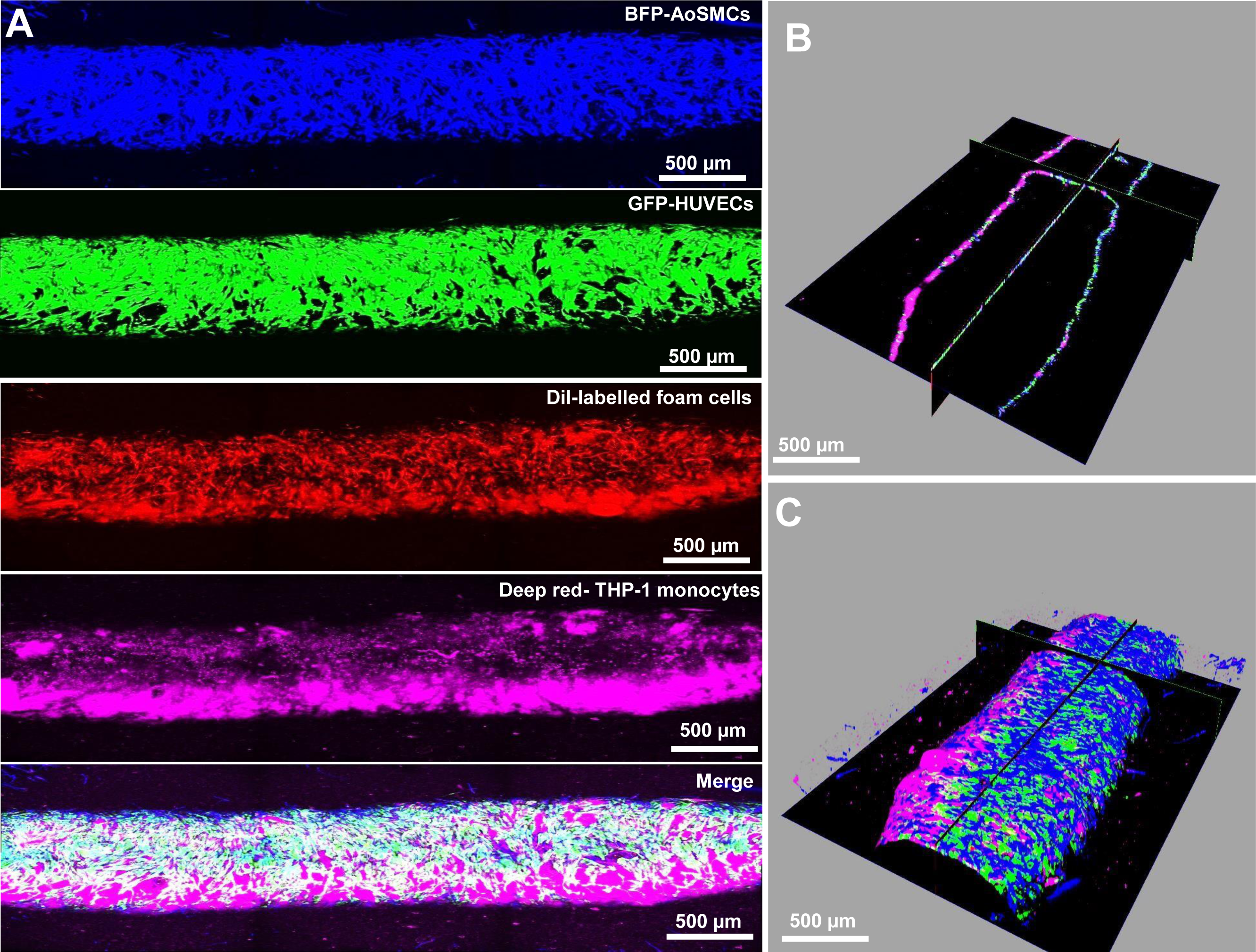
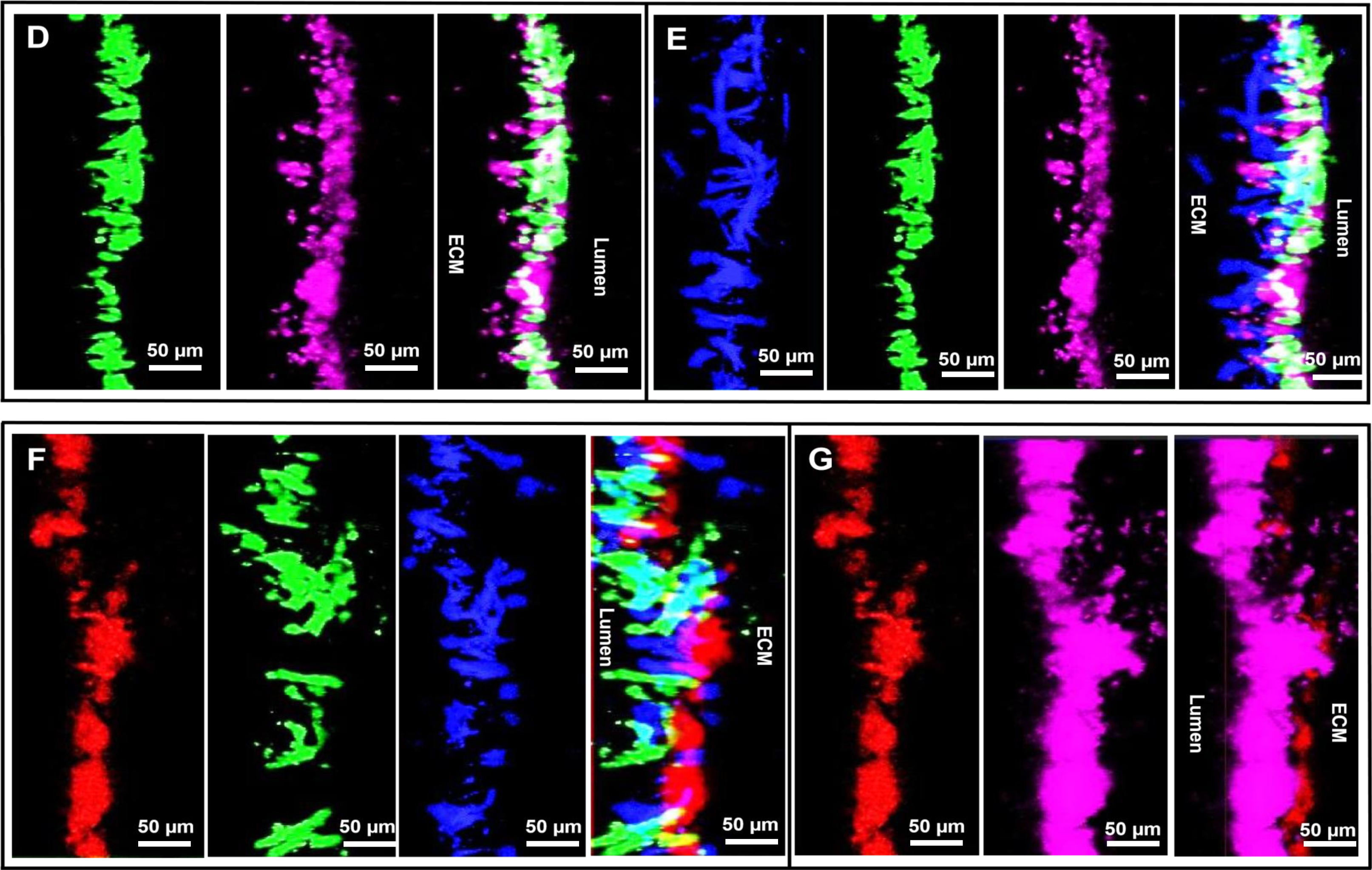
Human atherosclerotic vessel perfused for 24 hours with circulating monocytes in the microfluidic system. (A) Confocal micrograph shows the preserved endothelial monolayer formed by GFP-HUVECs (second panel, in green) surrounded by preserved BFP-AoSMCs (upper panel, in blue), and accumulated foam cells (third panel in red), with extravasated deep red stained THP-1 monocytes (fourth panel in magenta) and the merged image (lower panel) after 5 days of co-culture, from which 4 days static and 1 day perfused with circulating monocytes at a flow rate of 40µl/min in a human atherosclerotic vessel. (B) Composite display of the longitudinal and orthogonal cross section micrographs. (C) 3D reconstruction of half of a wall of the perfused macrovessel. (D) High magnification of a selected area in the longitudinal cross section with separated color channels show GFP-HUVECs (green), recruited circulating THP-1 cells (magenta), and composite image. Similar type of panels (E) showing BFP-AoSMCs (blue), GFP-HUVECs (green), recruited circulating THP-1 cells (magenta), and composite image, (F) Dil-oxLDL foam cells (red), GFP-HUVECs (green), BFP-AoSMCs (blue), and composite image, (G) Dil-oxLDL foam cells (red), recruited circulating THP-1 cells (magenta), and composite image.

Circulating THP-1 cells were also recruited to the sub-endothelial space, showing colocalization with VSMCs (Fig. 6e). The Dil-oxLDL signal indicated that most of the foam cells remained in the sub-endothelial space, co-localizing with the VSMCs (Fig. 6f). Recruited THP-1 cells also colocalized partly with Dil-oxLDL loaded foam cells, implying THP-1/foam cell interaction (Fig. 6g). In line with these findings, quantification of the composite images showed no significant change in BFP-hAoSMCs^+^ and GFP-HUVECs^+^ areas when compared to static atherosclerotic vessels, or atherosclerotic vessels perfused without circulating cells (Supplemental Fig. 4c, d). No difference was observed between THP-1 cell perfused atherosclerotic vessels when compared to THP-1 perfused control vessels (Supplemental Fig. 4i and j).

### Difference in response of vascular smooth muscle cells in static and perfused healthy and diseased vessels

VSMCs respond to biomechanical and inflammation signals in the atherosclerotic plaque and can switch between a quiescent contractile state to a migratory, proliferative synthetic phenotype^25–27^. To evaluate the impact of flow with and without circulating cells on the migratory response of VSMCs in the (atherosclerotic) vessels, accumulation of VSMCs in the area between the lumen and ECM borders were analysed using confocal maximum projection images. In healthy control vessels, 48h of flow at 40 µl/min increased the overall BFP-hAoSMC signal intensity (V/H-static-4d + flow, orange line) compared to static controls (V/H-static-4d, grey line) based on the peri-luminal distribution (Fig. 7a). However, segmentation of the periluminal space (into regions of 0-100, 100-200, and 200-300 micron away from the lumen) followed by quantification of the BFP-hAoSMC+ area in each segment, showed no significant changes in healthy control vessels in response to flow (Fig.7b). Likewise, no significant change in delta BFP-hAoSMC+ area (signal^+^ area in the 200-300 micron region subtracted from the signal^+^ area in the 0-100 micron region (delta segments)) was observed, indicating that flow did not alter VSMCs distribution (Fig. 7c).

**Figure 7.**
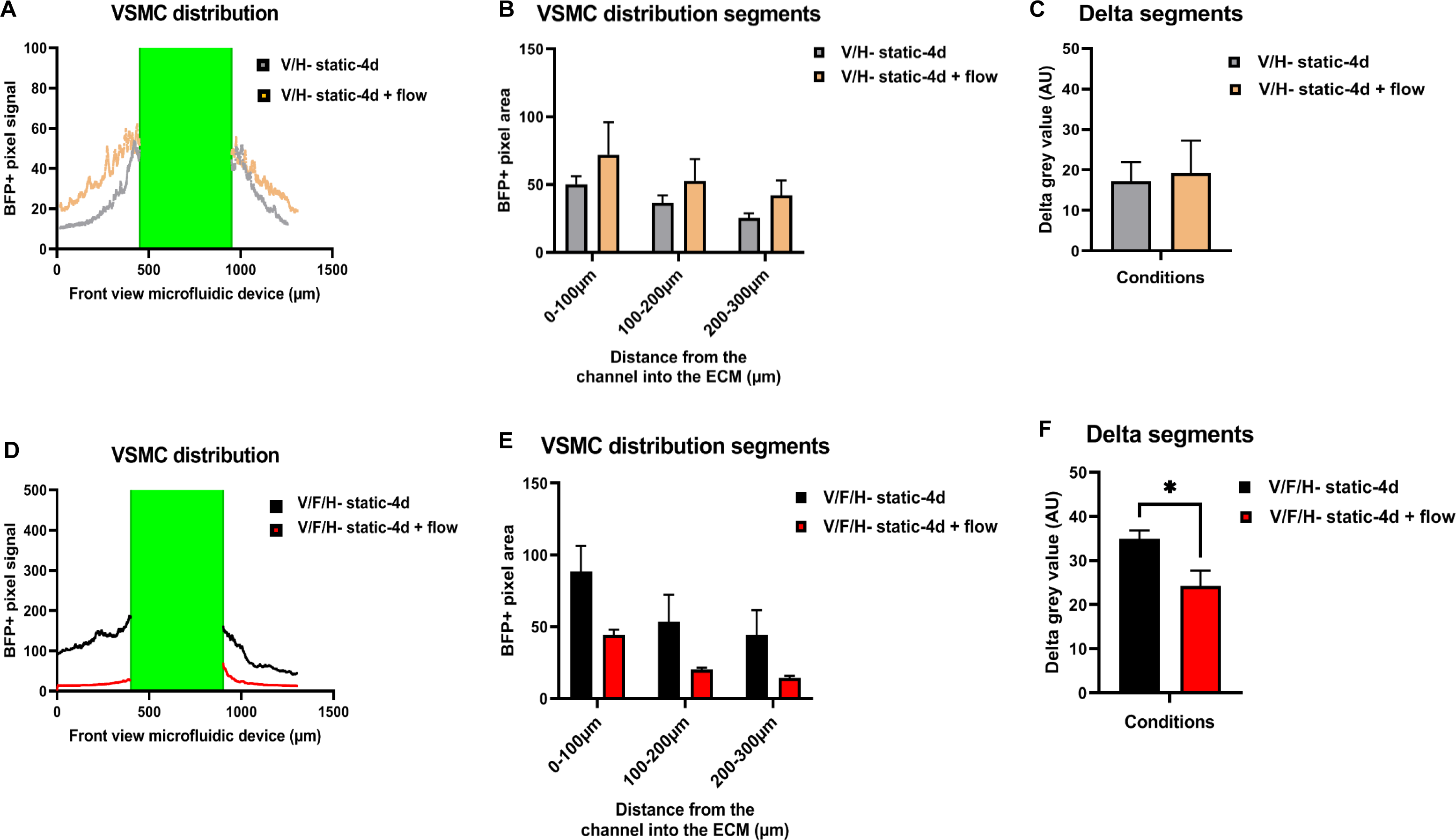
Impact of flow on VSMC distribution in healthy and atherosclerotic vessels. Impact of flow on VSMC distribution in healthy control (V/H) and atherosclerotic (V/F/H) vessels. (A) Representative view of BFP-VSMC distribution in a static and a perfused healthy control vessel. Distribution of average BFP signal intensity (grey value) over the cross-sectional vessel wall area of static control vessels (grey line) and control vessels perfused for 48 h (orange line). The green box marks the lumen area. (B) BFP-VSMC^+^ area distribution over predefined cross-sectional wall segments ranging 0-100, 100-200, 200-300 micron from the lumen, of static control vessels (grey bars) and control vessels perfused for 48 h (orange bars), N=3. (C) Bar graph showing the delta segments value of static and perfused control vessels, which was calculated by subtracting BFP-VSMC^+^ area values of segment 200-300 micron from segment 0-100 micron, N=3. (D) Representative view of BFP-VSMC distribution in a static and a perfused atherosclerotic vessel. Distribution of average BFP signal intensity over the cross-sectional vessel wall area of static atherosclerotic vessels (black line) and atherosclerotic vessels perfused for 24 h (red line). The green box marks the lumen area. (E) BFP-VSMC^+^ area distribution over predefined cross-sectional wall segments ranging 0-100, 100-200, 200-300 micron from the lumen, of static (black bars) and perfused atherosclerotic vessels for 24 h (red bars), N=4. (F) Bar graph showing the delta segments value of static and perfused atherosclerotic vessels, which was calculated by subtracting BFP-VSMC^+^ area values of segment 200-300 micron from segment 0-100 micron, N=4, *\*P<0.05*.

In atherosclerotic vessels, flow decreased the overall BFP-hAoSMC+ signal intensity (V/F/H-static-4d + flow, red line) compared to static controls (V/F/H-static-4d, black line) (Fig. 7d). Quantification of the GFP+ VSMC area in each segment, showed no significant changes in atherosclerotic vessels in response to flow (Fig.7e). However, a significant decrease in delta BFP-hAoSMC+ area (delta segments) was observed, indicating that flow promoted accumulation of VSMCs away from the lumen (Fig. 7f). To evaluate the impact of foam cells on VSMC distribution, atherosclerotic vessels were compared to control vessels after 4 days of static culture. A non-significant general increase in BFP-hAoSMC^+^ area was observed over all segments in foam cells loaded (atherosclerotic) vessels versus healthy controls without foam cell loading (Supplemental Fig. 5a). A significant increase in the delta BFP-hAoSMC+area was observed in the static atherosclerotic vessels versus control vessels under static conditions (Supplemental Fig. 5b). These data clearly demonstrate that presence of foam cells impacts VSMC distribution in the ECM. Similar results were observed when vessels were perfused with THP-1 cells: VSMC distribution away from the lumen was significantly decreased in atherosclerotic vessels (V/F/H-static-4d + THP-1 flow) compared to control vessels (delta segments of atherosclerotic vessels < healthy controls, Fig. 8a). Flowing with THP-1 cells significantly reduced the BFP-hAoSMC+ area in all segments in atherosclerotic vessels compared to control vessels (Fig. 8b).

**Figure 8.**
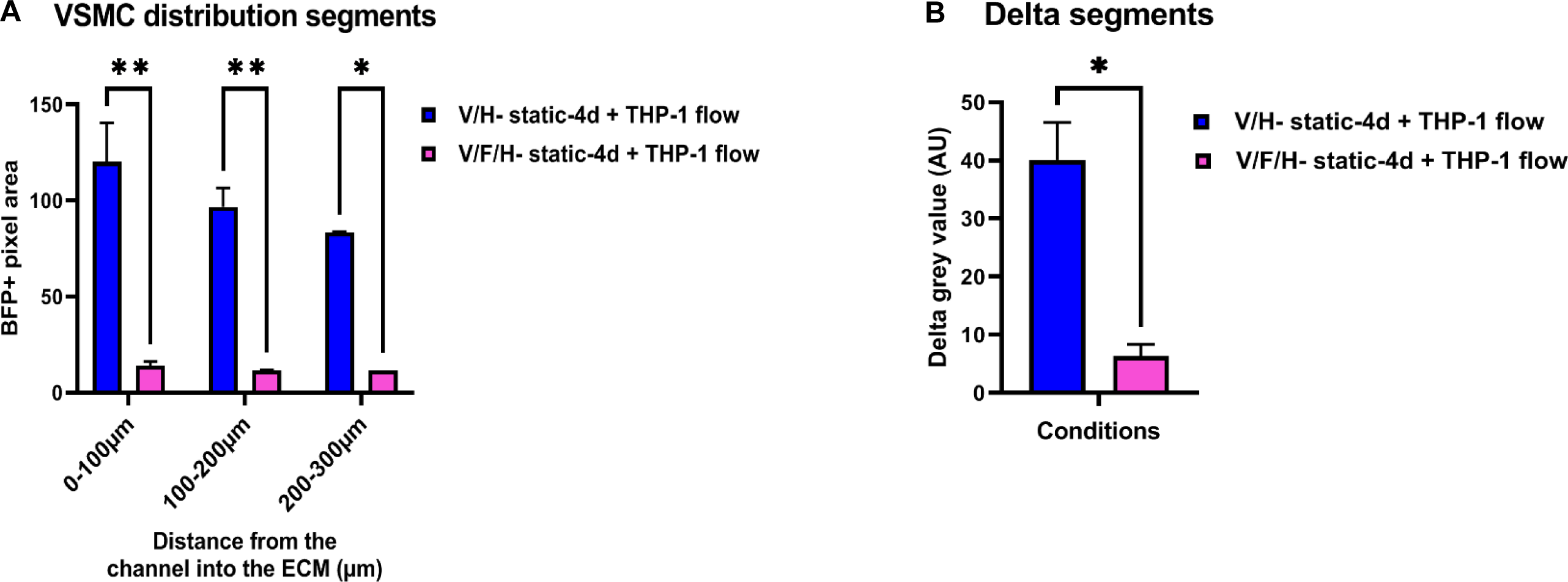
Impact of circulatory monocytes on VSMC distribution in healthy and atherosclerotic vessels. (A) BFP-VSMC^+^ area distribution over predefined cross-sectional wall segments ranging 0-100, 100-200, 200-300 micron from the lumen, of control (blue bars) and atherosclerotic vessels (red bars), perfused for 24 h with THP1 monocytes, N=3, *\*\*P<0.01, *P<0.05*. (B) Bar graph showing the delta segments value of control and atherosclerotic vessels, perfused for 24 h with THP-1 monocytes, calculated by subtracting BFP-VSMC^+^ area values of segment 200-300 micron from segment 0-100 micron, N=3, *\*P<0.05*.

### Enhanced recruitment of circulating monocytes in atherosclerotic versus control vessels

In response to *in vivo* inflammatory signals, circulatory monocytes migrate into the vessel wall and merge with the foam cell core during atherosclerosis progression. To evaluate if the model can mimic this process, we next evaluated total THP-1 monocyte area and their distribution in the lumen (free flowing monocytes) and sub-endothelial space (extravasated monocytes) of perfused healthy control (without foam cells) and atherosclerotic (with foam cells) vessels. Quantification of the deep red area showed significant increase in the total area of THP-1 monocytes in the atherosclerotic vessels with foam cells perfused for 24 h (pink bar) compared to control (blue bar, Fig. 9a), indicating that more THP-1 cells were present in the analysed atherosclerotic versus control vessels. Significantly higher significant grey values of the THP-1 deep-red signal were detected in the sub-endothelial space of atherosclerotic versus control vessels indicating that more THP-1 cells were recruited into the vessel wall (Fig. 9b). At the same time, no significant difference in free flowing THP-1 cells was found between atherosclerotic and control vessels (Fig. 9c) These data indicate that our microfluidic system can indeed successfully mimic the events of circulating cell recruitment in human atherosclerotic vessels.

**Figure 9.**
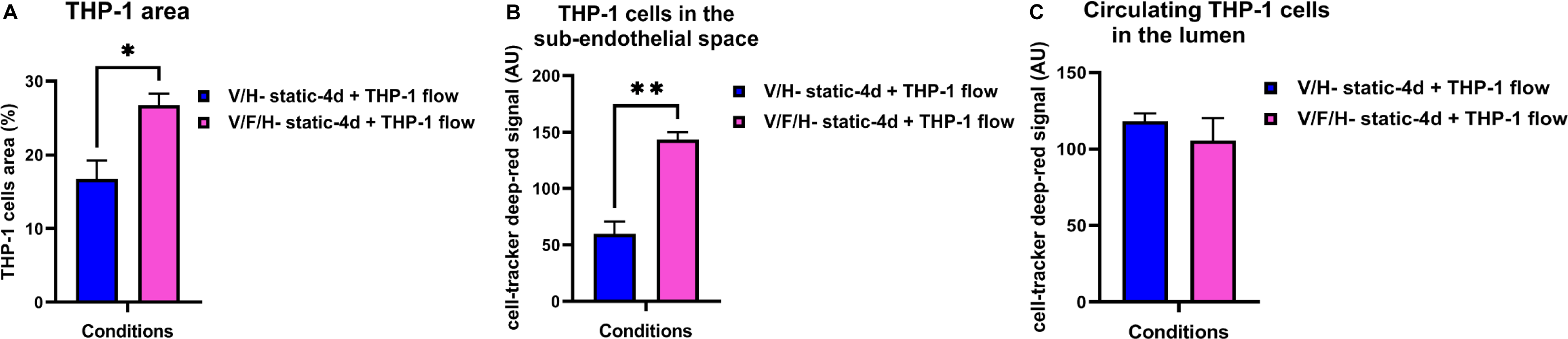
Distribution of circulatory monocytes in healthy and atherosclerotic vessels. (A) Bar graph presenting the total area of recruited deep-red labelled circulating THP1 monocytes in the atherosclerotic vessels (pink bar) compared to the control vessels (blue bar), after flowing for 24 h with THP-1 cells, N=3, *\*P<0.05*. (B) Average quantified deep-red signal intensity (grey value) over the sub-endothelial area of the cross-sectional vessel wall, in control and atherosclerotic vessels perfused with THP-1 cells, N=3, *\*\*P<0.05*. (C) Average quantified deep-red signal intensity (grey value) over luminal area, in control and atherosclerotic vessels perfused with THP-1 cells, N=3.

### Adaptation in gene expression of atherosclerotic vessels in response to flow and circulating monocytes

To evaluate the impact of flow, foam cells and circulating monocytes on the mRNA expression level of known atherogenic lipid, inflammation and adhesion markers, microfluidic channels were excised from the ECM gel after the last confocal readout. The qPCR analysis of the macrovessels showed significant upregulation of Low density lipoprotein-receptor, (LDLR) (a receptor that binds to Ox-LDL particles) under static culture in atherosclerotic versus control vessels (Fig. 10a). LDLR expression significantly decreased under flow compared to static in atherosclerotic vessels, whereas no flow response was observed in control vessels (Fig. 10b). The introduction of circulating monocytes did not impact LDLR expression in both atherosclerotic and control vessels (Fig. 10c).

**Figure 10.**
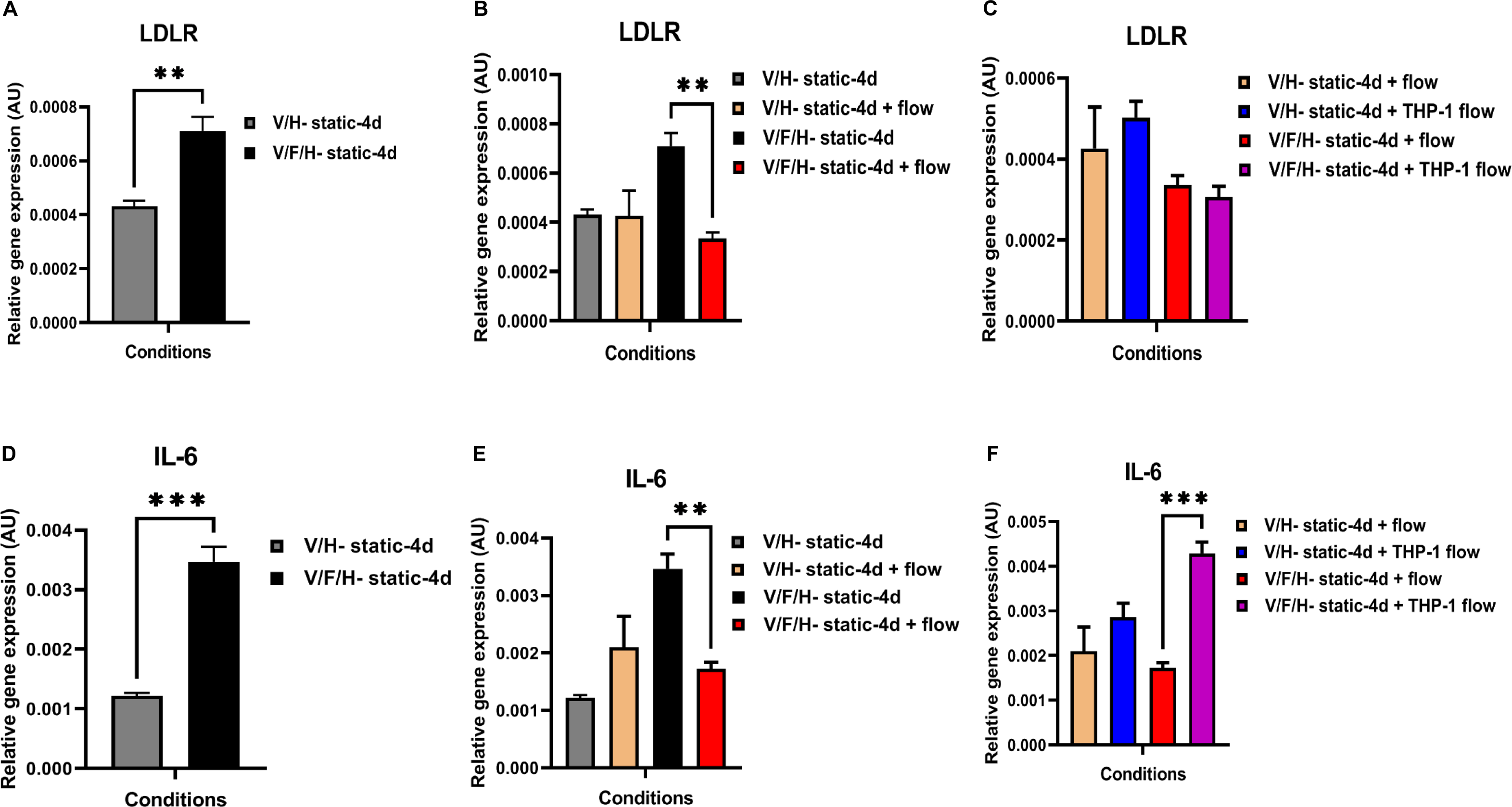
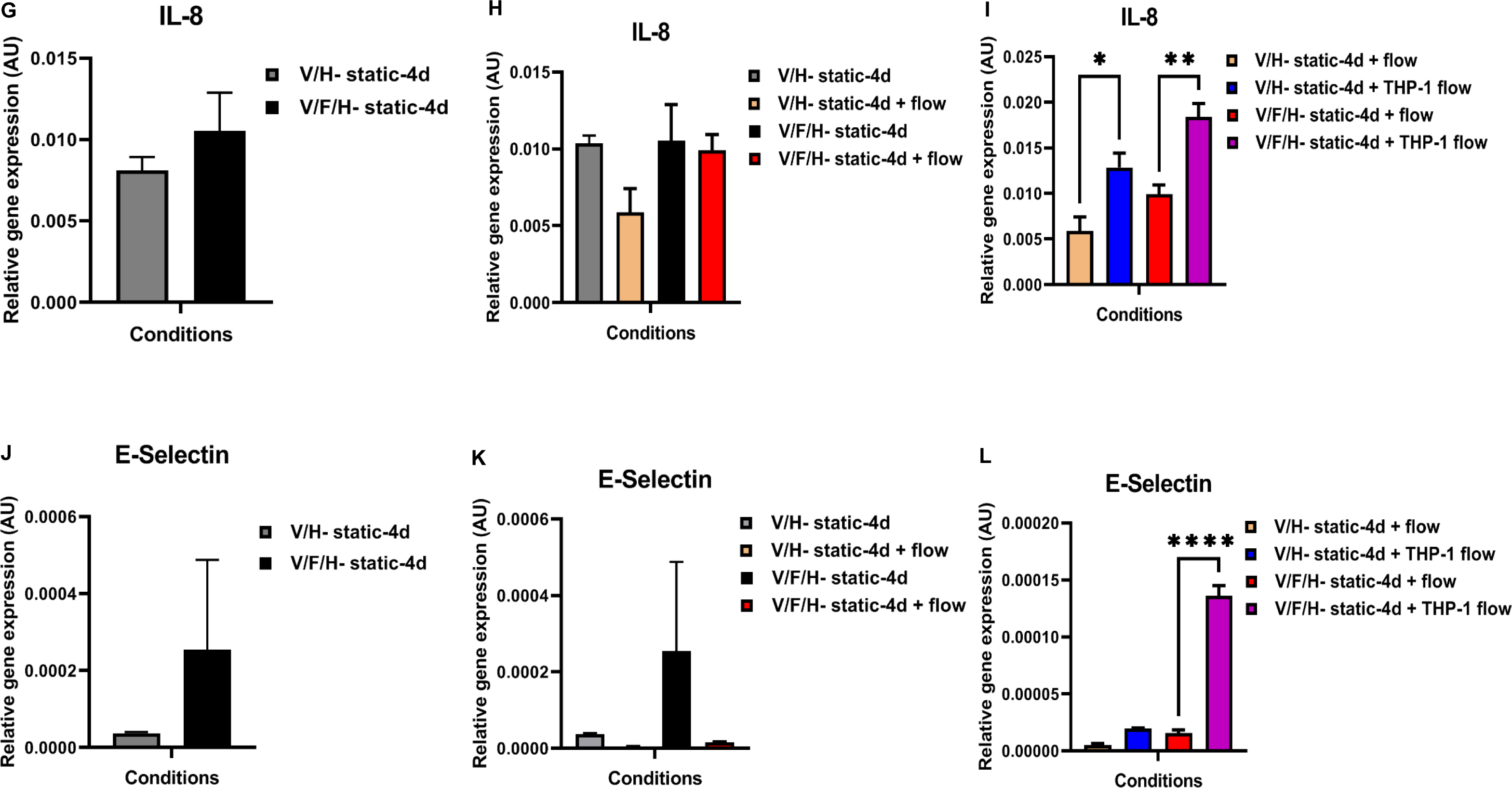
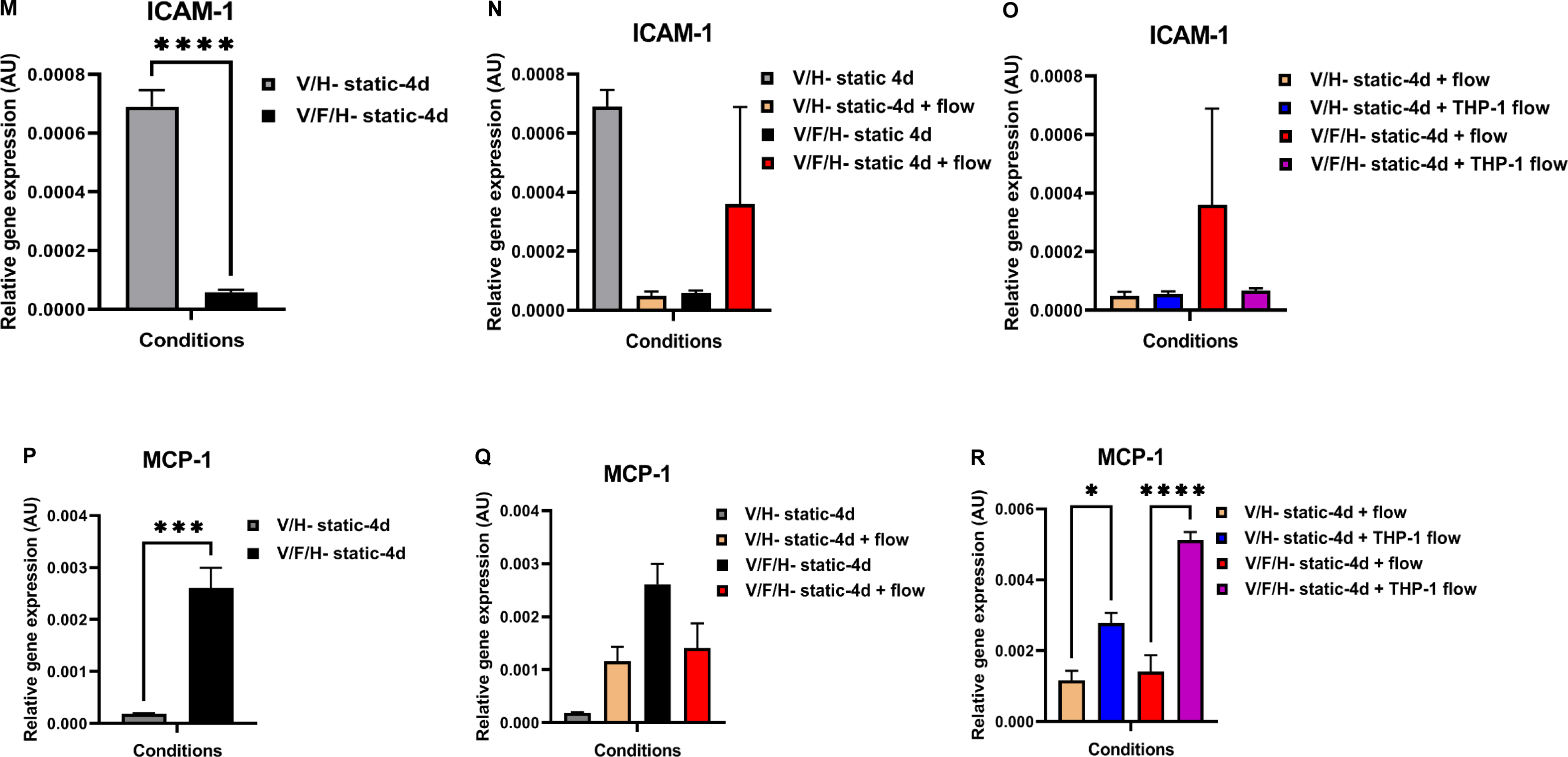
Effect of flow and circulating monocytes on foam cell, inflammatory and adhesion molecule marker expression. Relative mRNA levels expression of (A-C) LDLR, (D-F) IL6, (G-I) IL8, (J-L) E-selectin, (M-O) ICAM1, and (P-R) MCP1 in comparison between (A, D, G, J, M, P) control (grey bar) and atherosclerotic vessels (black bar) under static co-culture, (B, E, H, K, N, Q) in comparison between control and atherosclerotic vessels with and without flow, (C, F, I, L, O, R) in comparison between perfused control and atherosclerotic vessels with and without circulating THP-1 cells, N=4, *\*P<0.05*, *\*\*P<0.01*, \*\*\**P* <0.001, ****P<0.0001.

Next, expression levels of inflammatory cytokines Interleukin-6 (IL-6) and Interleukin-8 (IL-8) where analysed. IL-6 but not IL-8 mRNA levels were upregulated under static conditions in atherosclerotic versus control vessels (Fig. 10d, g) Flow downregulated IL-6 but not IL-8 in atherosclerotic vessels and did not affect IL-6 expression in control vessels (Fig. 10e, h).

When circulating monocytes were incorporated in the system, both, IL-6 and IL-8 mRNA levels were significantly upregulated in the atherosclerotic vessels. IL-8 mRNA was also increased in control vessels with perfused THP-1 cells (Fig. 10f, i). Adhesion molecules E-selectin and Intercellular adhesion molecule-1 (ICAM-1) were also analysed. Under static conditions, ICAM-1 but not E-selectin was significantly downregulated in the atherosclerotic vessels versus healthy controls (Fig. 10j, m). Flow had no significant impact on expression levels of both E-selectin and ICAM-1 (Fig. 10k, n). Perfusion with circulating THP-1 cells significant upregulated of E selectin expression in atherosclerotic vessels (Fig. 10l) but had no impact on ICAM-1 expression (Fig. 10o). Expression of monocyte chemoattractant protein-1 (MCP-1), a cytokine involved in monocyte recruitment, was significantly upregulated in static atherosclerotic versus control vessels (Fig. 10p). Flow alone did not significantly affect MCP-1 expression in healthy and diseased vessels (Fig. 10q), whereas perfusion with circulating THP-1 cells significantly enhanced MCP1 expression in both atherosclerotic and control vessels (Fig. 10r).

## Discussion

The aim of the study was to create a complex, perfused human atherosclerosis-OAC model. Here we present, for the first time, a microfluidics-based system with human atherosclerotic vessel-like structures. The most important findings in the present study are that the new system offers: 1) Perfusable macrovessels with preservation of endothelial monolayer, VSMC layers, and open lumen structure after exposure to controlled, unidirectional, continuous flow, with and without circulating monocytes. 2) Introduction of ox-LDL loaded foam cells creates atherosclerotic vessels with a triple-layered lesion-like structure (endothelium, foam cells, VSMC media layer) and an open lumen that is similarly preserved under flow, with and without circulating monocytes. (3) These human (early) atherosclerotic vessels show adaptation to hemodynamic and circulating cell exposure, including distinct changes in expression of lipid-associated, inflammatory and adhesion molecule markers, reduced accumulation of VSMCs near the lumen area in response to flow and circulation cells, and more THP-1 cell recruitment in the sub-endothelial space, compared to healthy control vessels. Combined, these findings demonstrate that our human atherosclerosis-OAC platform can be used to study the complex mechanistic events in early-stage atherosclerosis. With chip creation based on a polydimethylsiloxane mould casting technique^20^, the system is suitable for upscaling and high efficiency & low-cost testing of drug targets in atherogenesis.

In the native atherosclerotic lesion, interactions between VSMCs, ECs, and foam cells are vital for disease onset and progression. Early lesions are characterized by the sub-endothelial accumulation of foam cells forming the intima, VSMCs migration into same region followed by VSMCs proliferation, and the activation of the endothelium that facilitates recruitment of circulating immune cells, while the endothelium/media bilayer structure of the vessel wall generally remains mostly intact^10, 28^. Exposure to (changes in) hemodynamical forces is crucial to provide a continuous trigger for the lesion’s adaptive response during growth, and is a critical force that governs lesion interaction with circulating cells, which subsequently leads to local cell recruitment, enhanced inflammation and further plaque expansion^28–30^. To generate an ideal atherosclerosis-OAC model, all these factors should be considered. Despite the recent advances in microfluidic technology, only a few atherosclerosis-OAC models have thus far been reported, with most of them showing a significant trade-off in their designs for the before mentioned important factors. Qu *et al.* presented a tune-able microfluidic stenosis model for the study of endothelium-leukocyte interactions during atherosclerosis, which lacked incorporation of VSMCs and foam cells^17^. Other studies presented systems based on VSMC and EC coculture that lacked foam cells incorporation^31, 32^. In contrast, our new atherosclerosis-OAC platform not only incorporates 3D co-culture of ECs, VSMCs and foam cells, but also uses a protocol that allows the cells to form a layered structure that mimics the endothelium and media, with ox-LDL loaded THP-1 cells located in the sub-endothelial space, thereby creating a more plaque-like organization. This 3D layered setup allows the study of cell migration behaviour towards typical plaque anatomical reference points (endothelium, media, foam cell core), and could help reveal interaction patterns between ECs, VSMCs, and (recruited) THP-1 cells in both static and dynamic conditions.

In a more advanced platform recently presented by Su *et al.*, co-cultures of VSMCs and ECs were set up in an adapted version of the tension surface-based ECM patterning design to obtain a layered structure. The vessel wall was mimicked by seeding VSMCs and ECM in separate square channels, with ECs on top of the VSMCs layer. Ox-LDL loading of the model was combined with the addition of THP-1 cells and cytokine stimulation, but no experiments were conducted with foam cells, controlled laminar flow or circulating immune cells^18^. Similarly, Gu *et al.* presented a 3D stretch design with the co-culture of ECs and VSMCs forming a layered vascular wall structure, and THP-1 monocytes seeded on top, followed by Ox-LDL loading. This platform was capable of testing cell response to stretch, but the structure was grown on a flat surface, lacking the tubular vessel geometry. Flow as well as circulating cell interaction experiments were also not performed in this design^16^.

Indeed, most of these advanced microfluidics systems use square channels or other geometries that deviate from the native tubular form to recreate the atherosclerotic vessel tissue^16, 18^. However, to be able to conduct physiologically relevant flow (and circulating cell) experiments, continuous laminar flow should be introduced in a round tube structure that allows a symmetrical distribution of circulating particles. This cannot be optimally achieved by e.g. using square tubes, which cause particle disturbances^33^. For our platform, we used an ECM casting technique to obtain round tubular channels for cell seeding, thereby re-creating (atherosclerotic) vessels with a native vessel-like geometry. Our study findings have demonstrated that controlled, continuous unidirectional flow of healthy control and atherosclerotic arteries in our microfluidic system was feasible up to 48 hours, as no detrimental impact on vessel integrity in response to flow was observed, and arterial layered structure and sub-endothelial location of foam cells were overall conserved. More importantly, unlike previously described atherosclerosis-OAC designs, in which THP-1 cells were added under static conditions^21^, we showed that flowing with circulating THP-1 cells was also feasible in this system. Completely in line with observations in natural human atherogenesis^24^, enhanced adhesion and recruitment of circulating THP-1 cells into the subendothelial space of atherosclerotic vessels was observed compared to control vessels.

Perfusion of the atherosclerotic versus healthy vessels with and without circulating cells in our system also led to changes in VSMC behavior. Previous findings in human atherosclerosis and murine models demonstrated that during atherogenesis, accumulation of ox-LDL in the sub-endothelial space and subsequent foam cell differentiation, is coincided by phenotypic switching of VSMCs from a quiescent, contractile state to a proliferative synthetic phenotype, with migration of synthetic VSMCs into the intima to advance plaque growth^27, 34^. VSMCs migration has been explored in the previously described channel system presented by Su *et al.,* where it was demonstrated that VSMCs recruitment towards the sub-endothelial ECM space was enhanced after ox-LDL and/or inflammatory cytokine loading under static conditions^18^. In line with these findings, we observed increased accumulation of VSMCs in the vessel segments closest to the sub-endothelium in atherosclerotic vessels compared to the controls after 4 days of static co-culture in our system. Su’s model could not test the impact of flow on VSMC behaviour^18^, but using our model we could demonstrate that flow reduced VSMCs accumulation in the direct sub-endothelial space of the atherosclerotic vessels to baseline values (comparable to control vessels), whereas VSMC distribution in control vessels was not affected by flow. It has been reported that flow-induced shear stress in direct exposure experiments suppresses VSMCs proliferation and migration behaviour compared to static conditions^35, 36^. In addition, studies using perfused EC and VSMCs co-cultures demonstrated that shear stress sensed by the endothelium promotes synthetic to contractile phenotype switching in the underlying VSMCs via paracrine interaction^37^ and inhibits (contractile phenotype-associated) migration^38^. The findings in our atherosclerosis-OAC model are in line with these previous reports, demonstrating that our system can be used to study complex shear-induced crosstalk mechanisms between vascular cells. In native vessels, atherogenesis typically occurs at vascular sites exposed to deviations in shear stress such as oscillatory shear, and *in vivo* evidence have also demonstrated that they can act as causative factors^39^. As our atherosclerosis-OAC model is pump regulated, the current setup can also be used to study intra-plaque cell behaviour and monocyte recruitment in response to unidirectional versus oscillatory shear stress.

Increased expression of atherogenic factors LDLR, IL-6 and MCP-1^40–42^ was observed in atherosclerotic vessels compared to control vessels in static conditions. This enhanced expression was suppressed (mainly for LDLR and IL-6) after exposure to flow. It has been reported that phenotypic switching from contractile to synthetic VSMCs promoted expression of atherogenic factors such as IL-6 and MCP-1 (reviewed in^43^). The decline in expression of atherogenic factors by the atherosclerotic vessels in our platform may be caused by the induction of a more contractile phenotype in perfused versus static conditions. *In vivo* and *in vitro* studies show that recruitment of circulating monocytes into the atherosclerotic vessel wall coincides with increased expression of atherogenic factors^40–43^. Similarly, perfusion with circulating monocytes in our microfluidic platform enhanced expression of atherogenic factors IL-6, IL-8, and MCP-1 as well as the expression of adhesion molecule E-selectin, in atherosclerotic vessels as compared to flowing without circulating cells (Fig. 10l). E-selectin is upregulated in ECs during inflammation responses and is essential for circulating leukocyte-endothelial interaction during extravasation, implying an active positive feedback on THP-1 cell recruitment in our atherosclerosis-OAC model^44^. Combined, these data indicates that our new atherosclerosis-OAC platform mimics complex flow and circulating monocytes related processes in human atherogenesis on a tissue anatomical, cell-cell interaction and cell functional level.

### Limitations of the study

Without a necrotic core, fibroblasts, and VSMC/fibrous cap, the atherosclerosis-OAC model presented in this study mostly mimics a lesion in the early stage of atherosclerosis. This may limit the suitability of this system to studies that focus on flow and circulating cell (monocytes) interaction with the early lesion environment. The current model also uses human primary vascular cells. It could be further adapted for use of vascular cells of different sources, including induced pluripotent stem cells (iPSC) derived vascular cells from CAD patients, opening the door for the use of this system in high throughput personalized drug screening.

## Conclusions

The current study demonstrates, to our knowledge for the first time, a fluorescent labelled, quadruple cell co-culture-based atherosclerosis-OAC microfluidic platform that mimics the complexity of the 3D tubular atherosclerotic vessel with layered media and endothelium structure and sub-endothelial foam cell accumulation, which can be exposed to controlled continuous flow with and without circulatory THP-1 monocytes. This new system provides an advanced platform for 3D studies of vascular cell/immune cell interactions in early-stage atherosclerosis. Furthermore, this novel microfluidics model provides live-confocal imaging with side-to-side comparison between healthy with atherosclerotic vessels. Engineered with human cells it offers a humanized platform for in-depth mechanistic studies. Further adaptations of the model may include incorporation of patient-derived cells of iPSC and/or peripheral blood mononuclear cells (PBMCs) origin. Integration of this new atherosclerosis-OAC with genetic and GWAS studies with established biobanks could be especially rewarding, when dedicated disease platforms for patient subpopulations are created for personalized drug screening.

## Acknowledgments

We would like to thank Gert-Jan Kremers for providing training on Leica-SP8 DLS confocal microscopy at the Erasmus Optical Imaging Centre (OIC) and Lau Blonden for producing BFP-lentiviral supernatants. This research was financially supported by EC RESCUE grant (no. 801540 to C.C) and the the Dutch CardioVascular Alliance (an initiative with support of the Dutch Heart Foundation) Grant 2020B008 RECONNEXT (to D.J.D., M.C.V. and C.C.). We gratefully acknowledge the Gravitation Program “Materials Driven Regeneration”, funded by the Netherlands Organization for Scientific Research (024.003.013).

## Author contributions

Conceptualization: C.G.M.V.D., M.M.B., D.J.D., M.C.V., C.C.

Methodology and optimization: R.M., C.G.M.V.D., M.M.B., M.M.K., C.C.

Bioreactors construction: C.G.M.V.D., I.C

Confocal imaging and data analysis: R.M., C.G.M.V.D.

Lentiviral constructs: C.G.M.V.D., M.M.B.

Writing original draft: R.M., C.G.M.V.D., and C.C.

Writing-review and editing: R.M., C.G.M.V.D., D.J.D., M.C.V., M.M.K., and C.C.

## Data availability

The datasets generated during and/or analysed during the current study are available from the corresponding author on a reasonable request.

